# Tetraspanin-enriched microdomains play an important role in pathogenesis in the protozoan parasite *Entamoeba histolytica*

**DOI:** 10.1101/2024.03.27.586913

**Authors:** Han Jiang, Herbert J. Santos, Tomoyoshi Nozaki

## Abstract

Tetraspanins (TSPANs) are a family of proteins highly conserved in all eukaryotes. Although protein-protein interactions of TSPANs have been well established in eukaryotes including parasitic protists, the role they play in parasitism and pathogenesis remains largely unknown. In this study, we characterized three representative members of TSPANs, TSPAN4, TSPAN12, and TSPAN13 from the human intestinal protozoan *Entamoeba histolytica*. Co-immunoprecipitation assays demonstrated that TSPAN4, TSPAN12 and TSPAN13 are reciprocally pulled down together with several other TSPAN-interacting proteins including TSPAN binding protein of 55kDa (TBP55) and interaptin. Blue native PAGE analysis showed that these TSPANs form several complexes of 120-250 kDa. Repression of *tspan12* and *tspan13* gene expression led to decreased secretion of cysteine proteases. Meanwhile, strains overexpressing HA-tagged TSPAN12 and TSPAN13 demonstrated reduced adhesion to collagen. Altogether, this study reveals that the TSPANs, especially TSPAN12 and TSPAN13, are engaged with complex protein-protein interactions and are involved in the pathogenicity-related biological functions such as protease secretion and adhesion, offering insights into the potential regulatory mechanisms of tetraspanins in protozoan parasites.

## Introduction

Tetraspanins (TSPANs) are proteins composed of four transmembrane domains (TMDs), a small and a large extracellular loop (SEL and LEL, respectively), and well conserved among eukaryotes. LEL contains highly conserved characteristic motifs such as cysteine-cysteine-glycine (CCG) motifs. TSPANs are involved in the coordination of intracellular and intercellular functions as signal transduction, pattern recognition, antigen presentation, T-cell proliferation, cell migration, membrane protein trafficking, and membrane compartmentalization [1–5]. Moreover, TSPANs were shown to be involved in infection by several pathogens such as the human immunodeficiency virus (HIV), hepatitis C virus (HCV), human papillomavirus (HPV), *Plasmodium spp.,* and *Mycobacterium tuberculosis* [6]. In helminths such as *Opisthorchis viverrine* [7] and *Schistosoma mansoni*, TSPANs play a notable role in the formation of teguments [8], making them potential targets for vaccine development [9]. Furthermore, in the unicellular parasite *Trichomonas vaginalis*, research has identified the TSPAN TvTsp1 as a component of exosomes [10]. Additionally, TvTsp6, found in the flagella, is implicated in sensory reception and aids in the parasite’s migration during infection [11].

TSPAN-enriched microdomains (TEMs) in cell membranes, also known as TSPAN clusters, are involved in organizing cellular processes like adhesion and signaling [12]. It has been shown in humans that TSPANs in TEMs bind to a diverse range of molecules, including integrins, lipids, and other TSPANs, and the interaction with integrins has been proven to be pivotal in the regulation of cell adhesion and migration, which are essential processes in immune response and wound healing [13,14]. Additionally, the ability of TSPANs to bind to other TSPANs suggests complex intra-TEM interactions, which are crucial for organizing membrane proteins and modulating signal transduction. Furthermore, their association with lipids points to a role in maintaining membrane integrity and dynamics [15,16]. These microdomains are also crucial in diseases like cancer and immune disorders, mediate critical cell-cell interactions and signaling pathways [17].

Amidst this backdrop, *Entamoeba histolytica* is an intestinal protozoan parasite that causes amebiasis, a neglected cause of morbidity and mortality in low- and medium-income countries. It remains to be a global health problem affecting 50 million patients and causing 100,000 deaths annually [18]. Disease transmission occurs via ingestion of the infectious cyst through fecally-contaminated food or water. Excystation of the cyst after ingestion produces the trophozoites which invade the intestinal epithelium [19]. The parasite exploits virulence-related mechanisms which enable the invasion of intestinal epithelial tissue and extraintestinal tissue. Symptoms can vary, ranging from diarrhea, colitis, and dysentery to liver abscess [20].

While the structures and roles of TSPANs in other model eukaryotes including humans are well established [12], they remain largely unknown in *E. histolytica*. Our previous in silico genome-wide survey identified a total of 17 TSPANs in *E. histolytica* [21], but no functional study nor TEMs composition have been investigated. In this study, we characterized the constituents of potential complexes within the TEMs of *E. histolytica*. Our findings prove that *E. histolytica* likewise harbors TEMs comprised of a repertoire of distinct, yet consistently present TSPANs and associated binding proteins, analogous to structures observed in other organisms. Our results also indicate multifaceted roles of TSPANs and *Eh*TEMs involving multivesicular bodies (MVBs), exosomes, the endoplasmic reticulum (ER), and the nucleus. Furthermore, we provide evidence linking TSPANs to virulence, notably adhesion and protease secretion and adhesion, which are hallmark processes in the pathogenesis of *E. histolytica*. Specifically, our data indicated a diminished adhesive capability upon independent overexpression of two specific TSPANs. Additionally, targeted gene knock-down of certain TSPANs revealed a concomitant reduction in cysteine protease secretion. Overall, this study provides new insights into TSPANs and putative binding proteins within *E. histolytica*’s TEM hinting at their pivotal roles in the regulation of virulence of this organism.

## Results

### Expression and localization of HA-tagged TSPAN4 in amoeba vesicles

Our previous genome-wide search identified 17 TSPANs in *E. histolytica* [21]. Among them, TSPAN4 (EHI_075690) was particularly remarkable as the gene was reported to be expressed at a higher level in non-pathogenic strain than its isogenic (i.e., with the identical genetic background) pathogenic strain and that the parasite line that overexpressed TSPAN4 produced smaller amebic liver abscesses in the hamster model [22]. This result indicates that TSPAN4 likely negatively regulates virulence of this parasite in vivo [22]. These observations prompted us to investigate and characterize amebic TSPANs with TSPAN4 being an initial target. To determine the localization of TSPAN4, we created amebic transformant strain that expresses TSPAN4 with the carboxyl terminus hemagglutinin (HA)-tag (TSPAN4-HA). Immunoblot analysis of TSPAN4-HA overexpressing strain with anti-HA antibody immunoblot confirmed expression of TSPAN4-HA as a single band corresponding to the predicted size of 27 kDa (Fig 1A). We analyzed the subcellular localization of TSPAN4-HA by immunofluorescence assay (IFA). Double staining IFA with anti-HA and Hoechst 33342 suggest that TSPAN4-HA does not localize at nuclear structures (Fig 1B, top panel). Double staining IFA with anti-HA antibody and anti-Hgl [galactose and N-acetyl-galactosamine (Gal/GalNAc) lectin heavy subunit] monoclonal antibody showed that the primary location of TSPAN4-HA is not the plasma membrane, but internal vesicles (colocalization coefficient 0.280) (Fig 1B, middle panel). However, double staining with anti-Vps26 antiserum indicated that TSPAN4 was not highly co-localized with the endosomal marker Vps26 (colocalization coefficient 0.188) (Fig 1B, bottom panel).

**Fig 1.**
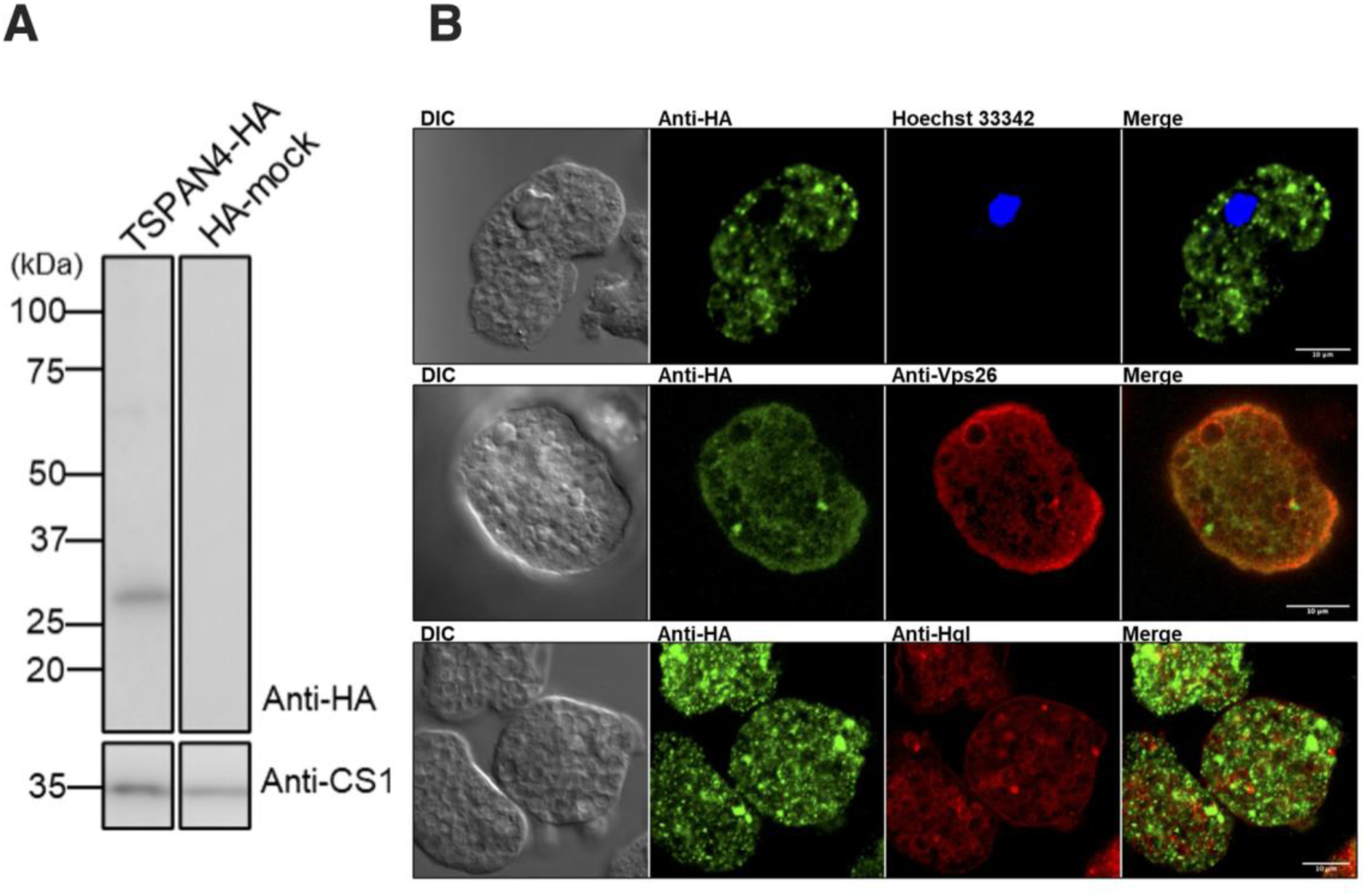
Expression and subcellular localization of TSPAN4-HA. (A) Immunoblot analysis of TSPAN4-HA in *E. histolytica* transformants. Approximately 20 µg of lysed cell samples from TSPAN4-HA-expressing transformant (TSPAN4-HA) and HA-mock-expressing transformant (HA-mock) were added to SDS-PAGE and immunoblot analysis using anti-HA antibody. CS1 (cysteine synthase 1) was detected by anti-CS1 antiserum as a loading control. (B) Representative immunofluorescence assay (IFA) micrographs of TSPAN4-HA double-stained with different antibodies. Top panel: mouse anti-HA (green) and DNA binding dye Hoechst 33342; middle panel: rabbit anti-HA (green) antibody and mouse anti-Hgl (red); bottom panel: mouse anti-HA (green) and rabbit anti-Vps26 (red) antiserum respectively. Scale bar, 10 µm.

### TSPAN4 interacts with other TSPANs and various other membrane proteins

To better understand the function of TSPAN4, we aimed to identify its interacting proteins by performing co-immunoprecipitation (co-IP) assay on TSPAN4-HA expressing trophozoites. Co-IP validation was performed by western blot analysis using anti-HA antibody. TSPAN4-HA was successfully precipitated and eluted (Fig 2A). The eluted fractions from the TSPAN4-HA and mock-HA co-IP samples were subjected to SDS-PAGE and silver staining (Fig 2B). The silver-stained gel showed unique bands detected in the TSPAN4-HA lane, specifically with an approximate molecular weight of 25 and 50 kDa, which were absent in the control mock-HA lane (Fig 2B, black arrows). The elution samples for both TSPAN4-HA and mock control were analyzed via mass-spectrometry (MS) to identify the binding partners of TSPAN4. Three independent co-IP experiments followed by MS analysis were performed to confirm reproducibility of the data.

**Fig 2.**
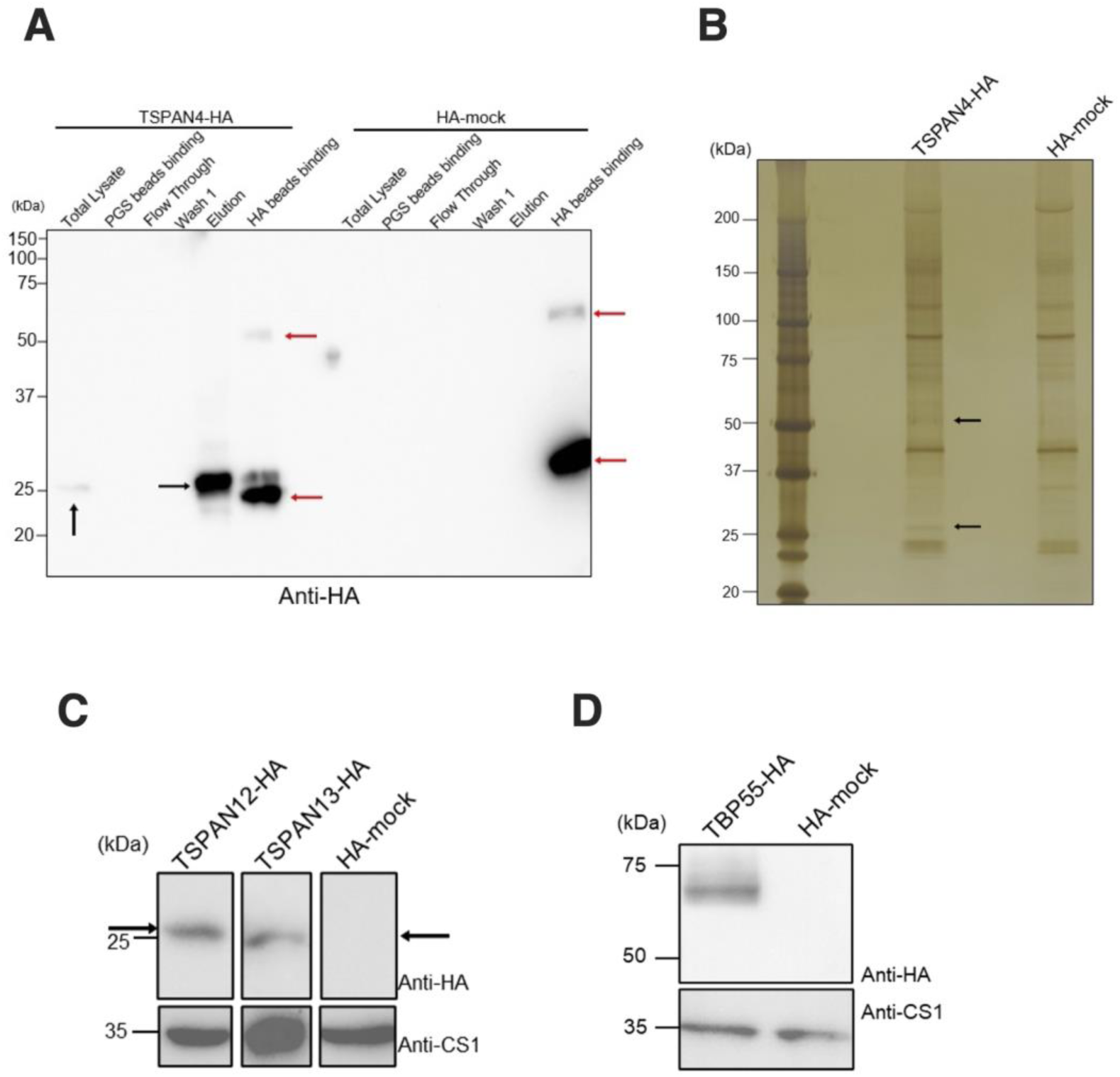
Co-immunoprecipitation assay of TSPAN4-HA. (A) Representative immunoblot analysis and silver staining of TSPAN4-HA in co-IP experiments. 5 µl of each fraction collected from co-IP experiments were added into SDS-PAGE and immunoblot analysis using anti-HA antiserum. Black arrows indicate TSPAN4-HA expression and red arrows indicate the heavy and light chain of anti-HA antiserum. (B) Silver staining was performed by adding 10 µl of eluted fraction of TSPAN4-HA and HA-mock in the co-IP. The black arrows suggest specific bands in the TSPAN4-HA sample. (C and D) Immunoblot analysis of TSPAN12-HA, TSPAN13-HA and TBP55 in *E. histolytica* transformants. Black arrows indicate TSPAN12-HA and TSPAN13-HA respectively.

To analyze the proteome data from the mass-spectrometry analysis, we prepared two lists of proteins labeled as “exclusive hits” and “enriched hits”. Exclusive hits indicate the protein candidates that were specifically pulled-down by HA-tagged TSPAN4 samples and not in the mock control. For the enriched hits, the ratio of quantitative value (QV) of normalized total spectra between HA-tagged TSPAN4 and mock control was computed and a threshold QV ratio value of 2.0 and above was set. A summarized table (S1 Table) was prepared, containing exclusive and enriched hits in all three independent co-IP trials, irrespective of QV, as well as proteins that have a mean QV over 5.0 in two out of three independent co-IP trials. The complete pull-down results are available as S1 File.

A total of four proteins were exclusively pulled down while one protein was enriched in TSPAN4-HA compared to mock-HA in all three trials (S1 Table). Moreover, TSPAN4 was validated to be exclusively precipitated in all three trials. Interestingly, two other TSPANs, TSPAN12 (EHI_091490) and TSPAN13 (EHI_107790) were likewise exclusively detected in all three trials with comparatively high mean QV of 13.6 and 19.0, respectively. This data indicates that in *E. histolytica* a similar nature of TSPANs forming complexes called TEM is conserved. In addition, two other proteins EHI_001100 and EHI_148910 were also pulled down in all three trials with the mean QV of 105.3 and 6.8, respectively.

### *E. histolytica* TSPAN4, TSPAN12, and TSPAN13 form heterogeneous tetraspanin-enriched microdomains

Upon identification of TSPAN12 and TSPAN13 as interacting proteins of TSPAN4-HA, we next constructed respective plasmids to express TSPAN12-HA and TSPAN13-HA in *E. histolytica* trophozoites. We confirmed the expression of TSPAN12-HA and TSPAN13-HA by anti-HA immunoblot analysis of corresponding cell lysates at approximately 28 kDa and 25 kDa, respectively (Fig 2C). To prove the interaction between these TSPANs, we performed reverse co-IP targeting TSPAN12-HA and TSPAN13-HA, followed by protein sequencing of eluted fractions by mass-spectrometry analysis. The co-IP and MS analyses were likewise independently repeated three times. Co–IP of TSPAN12-HA (S1A Fig) and TSPAN13-HA (S2A Fig) were validated as anti-HA immunoblotting revealed TSPAN12-HA and TSPAN13-HA bands appeared in respective eluted fractions. We also detected specific bands at around 25 and 50 kDa for TSPAN12 and 50 kDa for TSPAN13, which are supposed to be the potential binding partners of TSPAN12 and TSPAN13 in the silver-stained SDS-PAGE gels loaded with eluted fractions from mock control, TSPAN12-HA (S1B Fig) and TSPAN13-HA (S2B Fig). The results in the list for TSPAN12-HA (S2 Table) showed that the interacting proteins had a similar pattern with that of TSPAN4-HA. A total of seven proteins were pulled down by all three TSPAN12-HA co-IP trials. This includes TSPAN4, TSPAN12, TSPAN13, EHI_001100, and EHI_148910, which were also pulled down in all three trials of the TSPAN4-HA co-IP. Of note, EHI_001100 has a high QV compared with other binding partners of TSPAN4 and TSPAN12. This protein has been consistently identified in multiple independent co-IP experiments, indicating it is a binding partner of both TSPAN4 and TSPAN12. Hence, we named it TSPAN-binding protein of 55 kDa (TBP55). Another protein, EHI_148910, was detected with the mean QV of 6.8, 13.0, and 31.1 in all three trials of TSPAN4-HA, TSPAN12-HA and TSPAN13-HA, respectively. This protein was annotated as interaptin in AmoebaDB (https://amoebadb.org/), so we named it *Eh*interaptin.

On the other hand, the interactome of TSPAN13-HA showed a different pattern with that of TSPAN4 and TSPAN12. There are in total 57 proteins pulled down by all three trials of TSPAN13-HA co-IP, and neither TSPAN4 nor TSPAN12 were reversely pulled down by TSPAN13-HA (S3 Table). In addition, TBP55, although also pulled down in all three trials, only had a mean QV of 2.4 as compared to 105.3 and 65.5 in the TSPAN4-HA and TSPAN12-HA proteomic data, respectively.

### TBP55 is a key binding partner of amebic TSPANs

To further investigate the non-TSPAN binding partners of TSPAN4-HA and TSPAN12-HA, we established a TBP55-HA strain. Expression of TBP55-HA was successfully confirmed in the total lysate of *E. histolytica* (Fig 2D) by anti-HA immunoblotting. Co-IP of TBP55-HA was also performed three times (S3 Fig). The following MS analysis of TBP55-HA co-IP samples showed consistent pull down of TSPAN4, TSPAN12 and TSPAN13 in *E. histolytica* (Table 1). TBP55 also pulled down EHI_165070 at the mean QV of 17.1 in all three trials. EHI_165070 is annotated as short chain dehydrogenase, and was identified to be involved in fatty acid elongation [23]. Such binding suggests that TBP55 may associate with fatty acid elongation. Another enzyme EHI_076870, annotated as steroid 5-alpha reductase, was hit by TBP55-HA co-IP twice with a mean QV of 54.9. Steroid 5-alpha reductase was associated with amebic cell motility [24]. EHI_164540, annotated as β-ketoacyl reductase, was also detected twice in the TBP55-HA co-IP proteome. These three membrane proteins are supposed to locate to the ER membrane, which may also be one localization of TBP55. Besides, G protein-coupled receptor 1 (EhGPCR1, EHI_025100) was detected twice in the proteome with a mean QV of 8.0. GPCRs are considered as conserved interacting partners of TEMs [25].

**Table 1.** Mass-spectrometry results of HA-tagged TBP55 co-immunoprecipitation. Co-IP assay followed by mass-spectrometry analysis were performed as described in Materials and methods. Frequency of identification indicates the frequency for one protein to be detected in an exclusive or enriched manner in three independent trials. Mean of quantification value suggests the mean of quantitative value (normalized total spectra) calculated by scaffold 5 software, the value outside the parenthesis stands for HA-tagged TBP55 sample while the value inside the parenthesis stands for mock control. The order is sorted by frequency of identification firstly, and the mean of quantitative value secondly.

| Accession number | Frequency of identification | Mean of quantitative value | Annotation |
| --- | --- | --- | --- |
| EHI_001100 | 3 | 267.8 (0.9) | TBP55 |
| EHI_107790 | 3 | 62.8 (0) | TSPAN13 |
| EHI_091490 | 3 | 17.7 (0) | TSPAN12 |
| EHI_165070 | 3 | 17.1 (4.3) | Short chain dehydrogenase |
| EHI_076870 | 2 | 54.9 (20.0) | steroid 5-alpha reductase |
| EHI_040700 | 2 | 24.0 (9.1) | Coatomer subunit gamma |
| EHI_054830 | 2 | 19.3 (6.0) | Calcium-transporting ATPase |
| EHI_163540 | 2 | 14.3 (2.9) | β-ketoacyl reductase |
| EHI_020300 | 2 | 13.6 (5.7) | Ribosomal protein L15 |
| EHI_075690 | 2 | 12.5 (0) | TSPAN4 |
| EHI_163240 | 2 | 9.7 (3.4) | phosphatidate cytidyltransferase |
| EHI_179320 | 2 | 9.1 (3.3) | Wntless-like transmembrane protein |
| EHI_011940 | 2 | 8.4 (2.2) | Dolichyl diphosphooligosaccharide protein glycotransferase |
| EHI_025100 | 2 | 8.0 (2.0) | G protein-coupled receptor, EhGPCR1 |
| EHI_096210 | 2 | 6.6 (2.2) | Coated vesicle membrane protein |
| EHI_042870 | 2 | 6.4 (0) | Cell surface protease gp63, putative |
| EHI_086380 | 2 | 6.2 (2.5) | Hypothetical protein |
| EHI_068200 | 2 | 5.5 (2.5) | 60S ribosomal protein L31, putative |
| EHI_044830 | 2 | 5.2 (2.3) | 60S ribosomal protein L39, putative |
| EHI_093790 | 2 | 5.0 (0) | Dolichyl-diphosphooligosaccharide--protein glycosyltransferase subunit 1 |

### Nuclear proteins were pulled down with TSPANs and *Eh*interaptin

Several nuclear pore complex (NPC) proteins were identified in pull-down assay of TSPANs and *Eh*interaptin. Nucleoporin210 (Nup210, EHI_183510) was bound by TSPAN4-HA with a comparatively high mean QV of 22.3 in two out of three trials (Table 2). It was also pulled down by TSPAN12-HA with a QV of 7.6 in one out of three trials (Table 2). Moreover, TSPAN13-HA also pulled down Nup210 with a high mean QV of 29.0 in all three trials (Table 2). Another NPC protein Nucleoporin 54 (Nup54, EHI_010010) was pulled down by TSPAN12-HA with a QV of 6.8 in one out of three trials (Table 2), and by TSPAN13-HA with a mean QV of 9.5 in all three trials (Table 2). The nuclear pore complex serves as a critical gateway for the transport of molecules between the nucleus and the cytoplasm, regulating the passage of proteins, RNA, and other macromolecules to maintain cellular functions [26]. Nup210 and Nup54 are key components of this intricate structure, playing distinct roles in mediating nucleocytoplasmic transport and contributing to the overall functionality of the nuclear pore complex [27,28]. Overall, the pulldown of various nuclear proteins in TSPANs and the robust binding between TSPANs and *Eh*interaptin indicate the potential nuclear functions for TSPANs, especially TSPAN13. *Eh*interaptin also pulled down many nuclear proteins such as nucleolar protein 56 (Nop56, EHI_122740) with mean QV of 17.7 in all two trials. Nup210 again was pulled down by *Eh*interaptin the but only in one trial with a QV of 18.7, centromere binding protein cbf5 (EHI_115300) with a mean QV of 6.8 in all two trials and several ribosomal proteins including 40S ribosomal protein S6 (EHI_000590) with QV of 18.9 and 60S ribosomal protein L16-B (EHI_177190) with QV of 5.2. Uniquely, when looking at the list of binding proteins of TSPAN13-HA, there are more nuclear proteins than the other two TSPANs and TBP55 (S3 Table). For instance, there are Nucleoporin 210 (Nup210, EHI_183510), two SMC domain-containing proteins (*Eh*interaptin, EHI_152940), Cell division cycle protein 48 (EHI_045120), and Nucleoporin 54 (Nup54, EHI_010010). Several nuclear pore complex (NPC) members identified in the pull-down using TSPAN13-HA as bait indicate that TSPAN13 may be involved directly or indirectly in nuclear transport. Using cNLS Mapper prediction, a tool that identifies importin α-dependent nuclear localization signals (NLSs), https://nls-mapper.iab.keio.ac.jp), TSPAN13 possesses a bipartite NLS signal from positions 107 to 140.

**Table 2.** Nuclear proteins pulled-down by amebic TSPANs and their binding proteins. Nucleus-localized proteins found in TSPANs and *Eh*interaptin’s respective co-IP proteomes. The values on the upper cell of TSPAN4-HA, TSPAN12-HA, TSPAN13-HA, *Eh*interaptin-HA indicate the number of independent trials the protein was pulled down by the respective bait. The values on the lower cell indicate mean QV; number outside parenthesis is QV in TSPANs or *Eh*interaptin, while number inside parenthesis is QV in the mock control. ND, no detection.

| Accession number | Annotation | TSPAN4-HA | TSPAN12-HA | TSPAN13-HA | *Eh*interaptin-HA |
| --- | --- | --- | --- | --- | --- |
| EHI_183510 | Nup210 | 2 out of 3 | 1 out of 3 | 3 out of 3 | 1 out of 2 |
|  |  | 22.3 (0) | 7.6 (0) | 29.0 (1.9) | 18.7 (3.2) |
| EHI_010010 | Nup54 | ND | 1 out of 3 | 3 out of 3 | 1 out of 2 |
|  |  |  | 6.8 (0) | 9.5 (3.5) | 4.9 (1.3) |
| EHI_122740 | Nop56 | 1 out of 3 | ND | ND | 1 out of 2 |
|  |  | 10.1 (4.8) |  |  | 17.7 (8.2) |
| EHI_115300 | Cbf5 | ND | 1 out of 3 | ND | 2 out of 2 |
|  |  |  | 14.1 (4.0) |  | 6.8 (1.9) |
| EHI_148910 | *Eh*interaptin | 3 out of 3 | 2 out of 3 | 3 out of 3 | 2 out of 2 |
|  |  | 6.8 (0) | 13.0 (1.1) | 31.1 (1.5) | 85.5 (1.9) |
| EHI_152940 | SMC-domain containing protein | 2 out of 2 | 2 out of 3 | 3 out of 3 | 1 out of 2 |
|  |  | 8.0 (0) | 4.2 (3.8) | 15.1 (1.5) | 56.2 (0) |

### Cellular localization of TSPAN12, TSPAN13, TBP55, and *Eh*interaptin

Although the subcellular localization of three TSPANs (TSPAN1, 2 and 4) in *E. histolytica* was described as mostly cytosolic vesicular compartments in our previous [21] and present studies, the subcellular localization of other TSPANs remains unknown. To further clarify the subcellular localization of TSPAN12, TSPAN13, and their binding partners, TBP55 and *Eh*interaptin, the IFAs were performed on the transformants that expressed TSPAN12-HA, TSPAN13-HA, TBP55-HA, and *Eh*interaptin-HA, respectively. The anti-HA signals of fixed TSPAN12-HA, TSPAN13-HA and TBP55-HA trophozoites were similar to the punctate patterns also observed in our TSPAN4-HA IFA. Double-staining IFA using anti-HA antibody and anti-binding immunoglobulin protein (BiP) antiserum (ER marker) showed partial co-localization of TSPAN12-HA (colocalization coefficient 0.465), TSPAN13-HA (colocalization coefficient 0.652), and TBP55-HA (colocalization coefficient 0.221), respectively, to the ER marker (Fig 3A). Meanwhile, *Eh*interaptin-HA showed obvious nuclear localization by Hoechst 33342 nuclear staining (colocalization coefficient 0.999, Fig 3B, top panel). However, TSPAN13-HA appears to be localized in close proximity to membrane structures surrounding Hoechst 33342 staining, as seen in the DIC image (Fig 3B, bottom panel), supporting the premise that TSPAN13 is potentially involved in nucleocytoplasmic transport.

**Fig 3.**
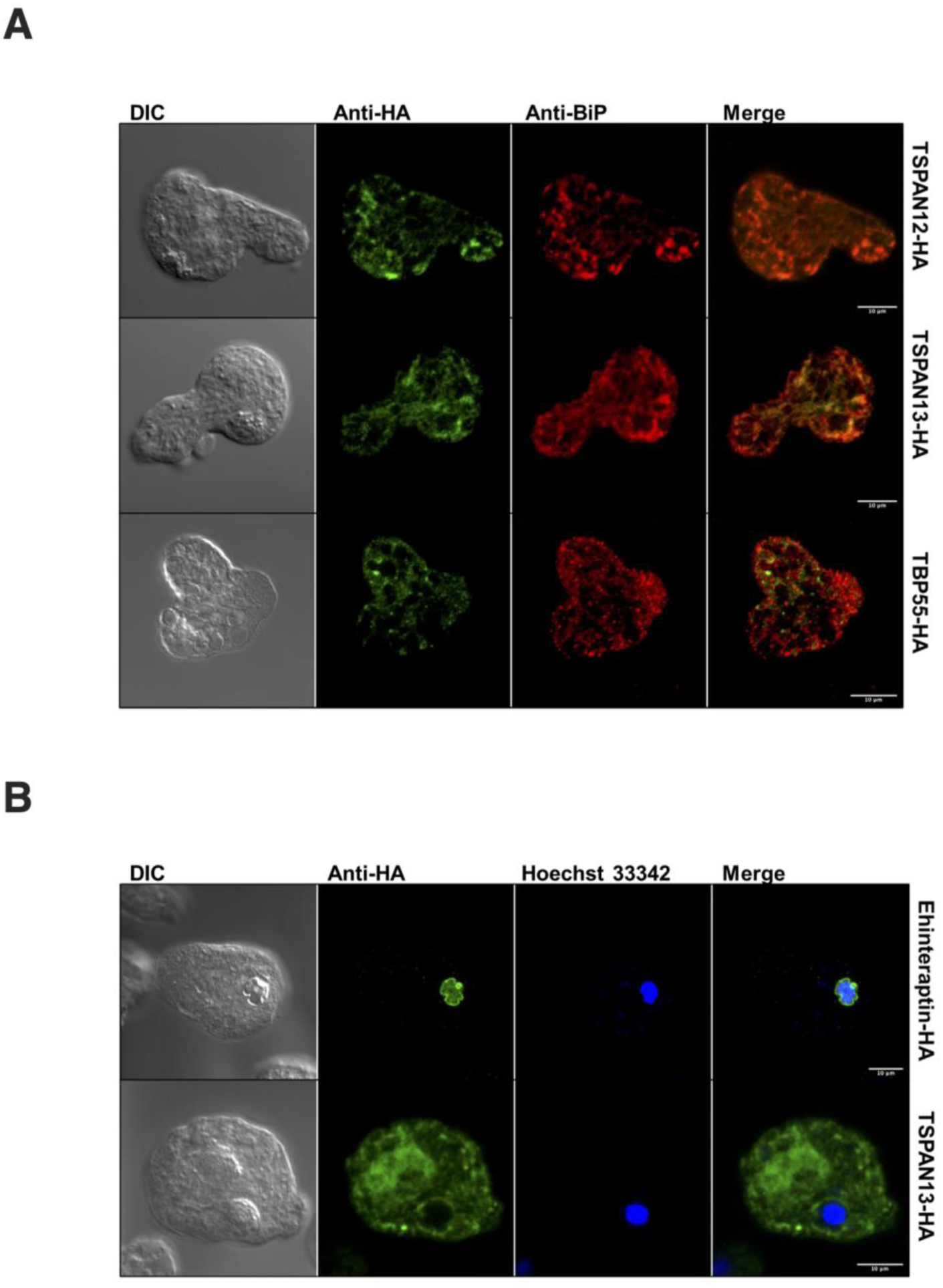
Subcellular localization of TSPAN12-HA, TSPAN13-HA, TBP55-HA and *Eh*interaptin-HA. (A) Representative immunofluorescence assay micrographs of TSPAN12-HA (top panel), TSPAN13-HA (middle panel) and TBP55-HA (bottom panel) double-stained with mouse anti-HA (green) and rabbit anti-BiP (red) respectively. Scale bar, 10 µm. (B) Representative immunofluorescence assay micrographs of *Eh*interaptin-HA and TSPAN13-HA double-stained with mouse anti-HA antibody (green) and DNA binding dye Hoechst 33342 (blue) respectively. Scale bar, 10 µm.

### Blue native-PAGE revealed heterogeneous complexes of TSPAN4-HA, TSPAN12-HA, and TBP55-HA

To investigate potential complex formation and determine the approximate size of the individual TSPAN-associated complexes at TEMs comprising of TSPAN4-HA, TSPAN12-HA, TSPAN13-HA, or TBP55-HA, we conducted organelle membrane fractionation followed by blue native (BN)-PAGE using the transformants of TSPAN4-HA, TSPAN12-HA, TSPAN13-HA, TBP55-HA, and mock-HA strains. Protein complexes were found in the membrane fraction of trophozoites expressing TSPAN4-HA, TSPAN12-HA, TBP55-HA, but not TSPAN13-HA. A complex of approximately 100 kDa containing TSPAN4-HA was identified based on immunoblot analysis of the BN-PAGE separated membrane fraction (Fig 4A, black arrow). On the other hand, four complex bands were detected after anti-HA immunoblotting of the BN-PAGE separated membrane fraction of TSPAN12-HA at around 250, 230, 160, and 100 kDa (Fig 4A, red arrows). Meanwhile, TBP55-HA was determined to form two complexes with approximate molecular weights of 250 and 230 kDa (Fig 4B, blue arrows). Vps26-1/2 of the retromer, which is involved in the retrograde transport of hydrolase receptor from the endosomes to the trans-Golgi network [29] and shown to form mono-to-tetrameric complex ranging 170-700 kDa, was used as a representative complex forming control protein (Watanabe et al., unpublished).

**Fig 4.**
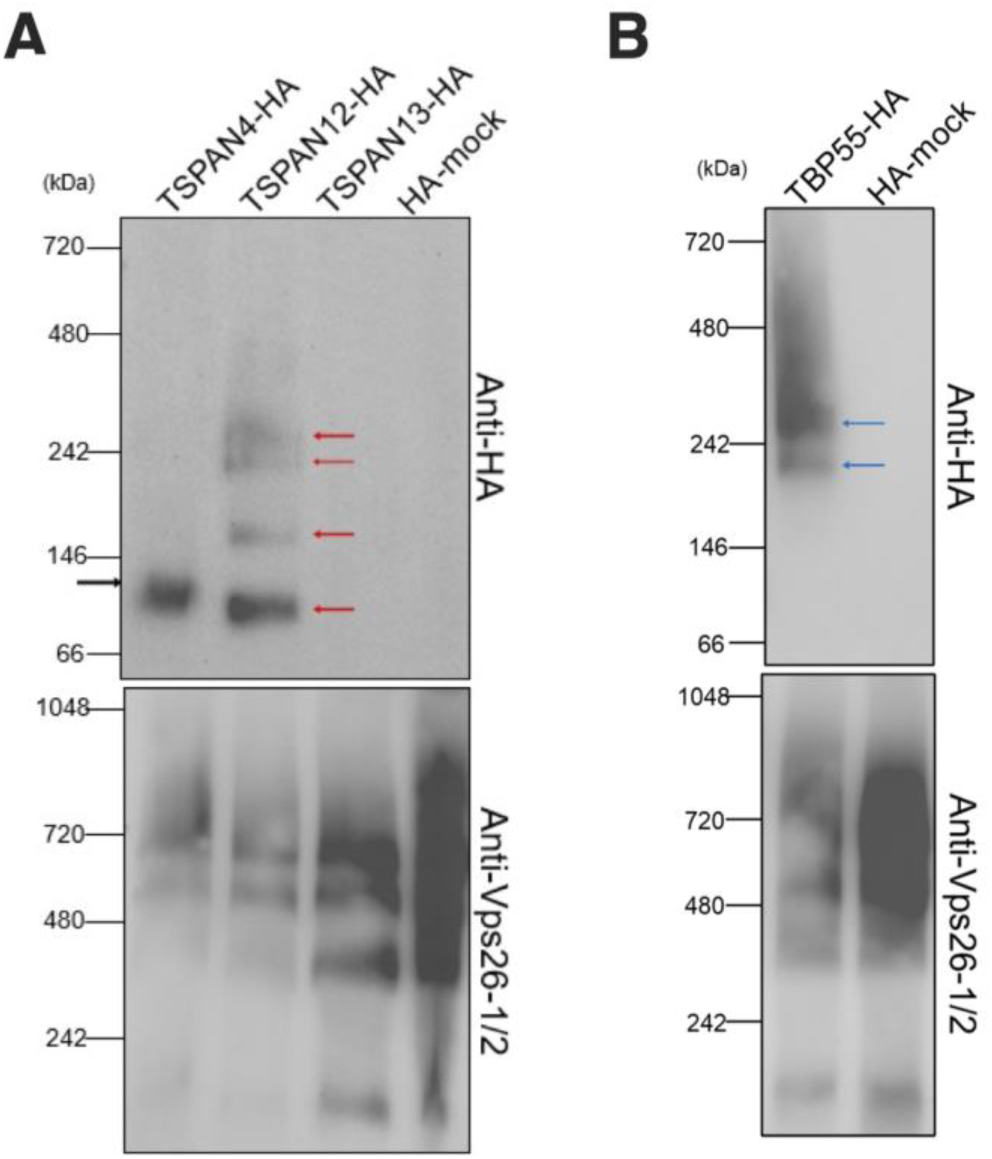
Blue native-PAGE indicates TSPAN4-HA, TSPAN12-HA and TBP55-HA forming complexes in membrane fraction. Transformants expressing (A)TSPAN4-HA, TSPAN12-HA, TSPAN13-HA and (B)TBP55-HA together with HA-mock control were used for membrane fractionation and BN-PAGE. Immunoblot analyses were performed by anti-HA, and anti-Vps26-1/2 using as loading control. Black arrow indicates protein complex involved TSPAN4-HA. Red arrows indicate protein complexes involving TSPAN12-HA. Blue arrows indicate protein complexes involving TBP55-HA.

### TSPANs are involved in both adhesion and cysteine protease secretion in *E. histolytica*

Regulation of cell adhesion is considered as one feature of TSPAN family proteins [30]. Thus, an in-vitro adhesion assay on collagen-coated plastic plate was performed on amebic lines in which each of *tspan4, tspan12, tspan13,* and *tbp55* genes was individually silenced. To this end, *E. histolytica* G3 strains transformed with psAP2-TSPAN4, psAP2-TSPAN12, psAP2-TSPAN13, and psAP2 (control) were used for gene silencing strain establishment [31]. The respective genes for *tspan4, tspan12, tspan13,* and *tbp55* were almost completely abolished based on the electrophoresis of PCR-amplification of corresponding transcripts (Fig 5A). The mean adhesive cell percentage of *tspan4*gs, *tspan12*gs, *tspan13*gs and psAP2 control transformants was 46.6% ± 5.3%, 50.6% ± 6.5%, 46.6% ± 6.1%, and 46.4% ± 4.2%, respectively (Fig 5B). The P-values of unpaired t-test of all the gs strains are over 0.05, suggesting that *tspan* gene silencing did not affect adherent ability with statistical significance.

**Fig 5.**
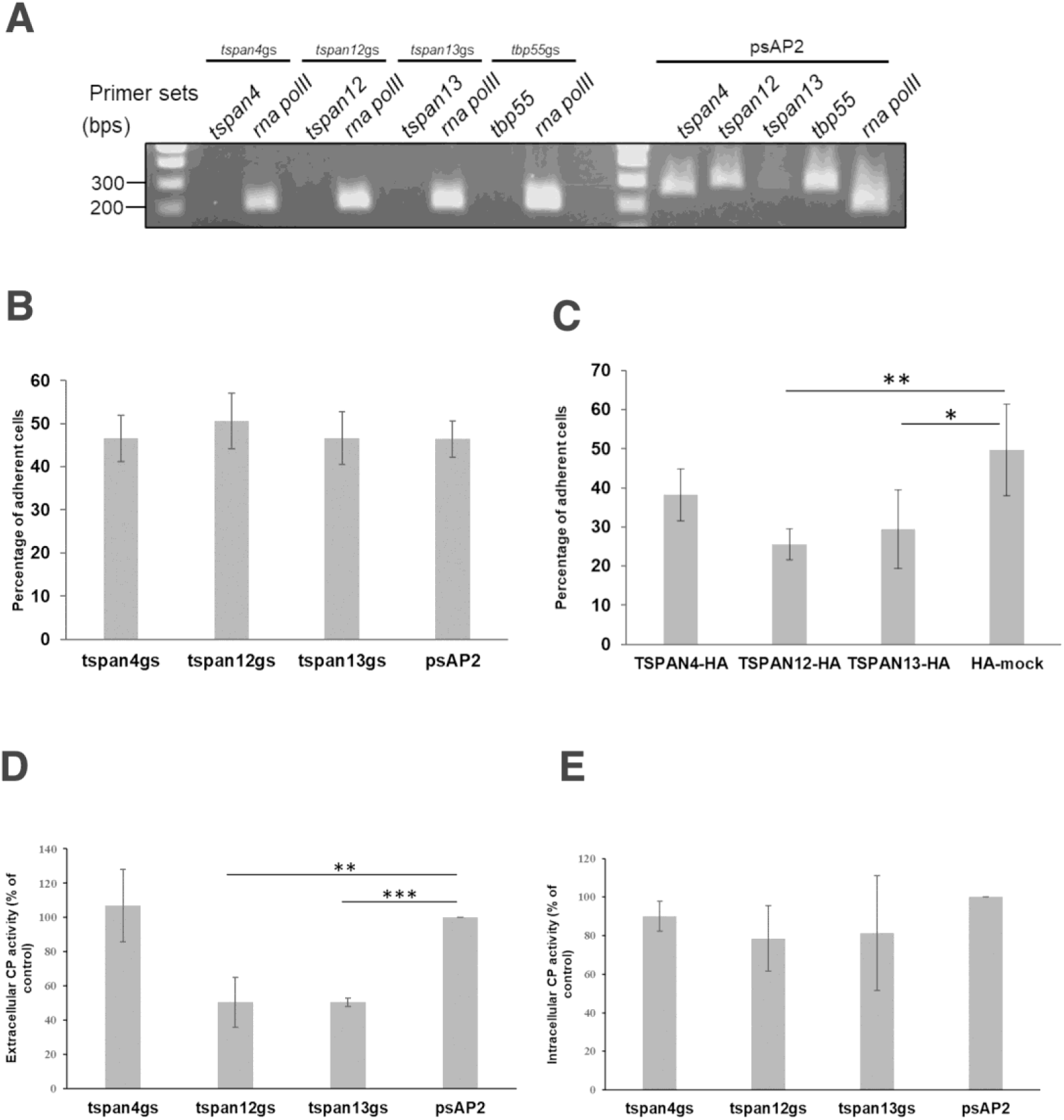
Tetraspanins variably affect the virulence of *E. histolytica*. (A) Confirmation of gene silencing by RT-PCR analysis of mock transfected and psAP2 derived gene silenced (GS) strains. Transcripts of *tspan4*, *tspan12*, *tspan13*, *tbp55* and *rna polymerase II* genes were amplified by RT-PCR from cDNA isolated from the transformants and examined by agarose gel electrophoresis. (B) Adhesion of CellTracker Green-stained *tspan4*, *tspan12* and *tspan13* gene silenced strains and control. Fluorescence intensity was detected before and after washing. The assay was performed five times independently. Data points in the graph show the mean and error bars represent standard deviation for five trials. Statistical significance was examined with Student’s unpaired t-test. (C) Transformants expressing TSPAN12-HA and TSPAN13-HA showed lower adhesive ability against HA-mock control strain. (** P < 0.01, p-value = 0.0078; * P<0.05, p-value = 0.0395). (D) *tspan*12 and *tspan*13 gene silenced strains secrete comparatively lower CP activity in extracellular environment, (E) While there is no significant difference in intracellular CP activity. Intracellular CP activity measuring based on cell lysate and extracellular CP activity measuring based on the supernatant after two centrifugations of 3,000 g and 15,000 g respectively. Samples were mixed with substrate z-Arg-Arg-MCA and underwent 60 minutes of fluorescence intensity detection. Data points in the graph show the mean and standard deviation for a total three independent trials with triplicates samples. Statistical significance was examined with Student’s unpaired t-test (*** P < 0.001, p-value = 0.0000043, ** P<0.01, p-value = 0.0042).

We also examined adhesive ability of transformants expressing TSPAN4-HA, TSPAN12-HA, TSPAN13-HA, and HA-mock. The experiments were repeated for four independent trials. The mean adhesive cell percentage of TSPAN4-HA, TSPAN12-HA, TSPAN13-HA, and HA-mock were measured at 38.2% ± 6.6%, 24.5% ± 4.0%, 29.5% ± 10.1% and 49.8% ± 11.7%, respectively (Fig 5C). The results suggested that TSPAN12-HA and TSPAN13-HA strains showed statistically significant lower adhesion ability (*p* = 0.0078, 0.0395, respectively) than the HA-mock control strain. Notably, TSPAN12-HA overexpressing trophozoites showed almost half the number of adhesive cells compared to HA-mock control.

To examine if TSPANs are involved in another major cellular activity pivotal to pathogenicity, cysteine protease (CP) activity was measured. CP assay was performed using trophozoites of *tspan4*gs, *tspan12*gs, *tspan13*gs, and psAP2 control strains. CP activities are shown as percentage relative to the vector control transformant (psAP2 control) (Fig 5D and 5E). We found approximately 50% reduction in the extracellular (secreted) CP activity in *tspan12*gs (50.6 ± 14.6%, *p* = 0.0042) and *tspan13*gs (50.4 ± 2.5%, *p* = 0.0000043) strains, compared to vector control, while no change was observed for *tspan4*gs strain (107.0 ± 21.3%, *p* = 0.6001) (Fig 5D). On the other hand, the intracellular CP activity did not vary significantly in all silenced strains, when compared against mock control. Measured intracellular CP activity of *tspan4*gs, *tspan12*gs, and *tspan13*gs strains were 90.0 ± 7.7%, 78.5 ± 17.0%, and 81.2 ± 29.9%, respectively, to that of the vector control (Fig 5E). Taken together, *tspan12* and *tspan13* gene silencing caused reduction in extracellular, but not intracellular, CP activity, consistent with the premise that TSPAN12 and TSPAN13 are involved in CP secretion.

## Discussion

TSPANs play a crucial role in diverse biological processes by engaging with various partners within microdomains [32]. In this study, we demonstrated the constituents and binding profiles of three TSPANs in the TEMs in *E. histolytica*. We further demonstrated that amebic TSPANs are involved in core virulence mechanisms of this parasite such as adhesion and CP secretion.

Based on multiple proteomic analyses, it is evident that TSPAN4 consistently binds with two other TSPANs, TSPAN12 and TSPAN13, as well as other proteins such as TBP55 and *Eh*interaptin. The interplay among different TSPANs is a common phenomenon conserved in eukaryotes [33]. Although there are 17 TSPANs in *E. histolytica*, the specific and exclusive interactions between TSPAN4, TSPAN12, and TSPAN13 stand out amidst the array of other TSPANs. Upon examination of the interacting partners of TSPAN12 and TSPAN13, distinct patterns emerged in comparison to that of TSPAN4. TSPAN12 efficiently bound TSPAN4, whereas TSPAN13 exhibited an inability to pull down both TSPAN4 and TSPAN12. This discrepancy suggests that TSPAN13 may not directly or strongly bind with TSPAN4 and TSPAN12. Additionally, the inability of TSPAN13-HA to pull down other TSPANs raises the possibility that the attached epitope tag could be a contributing factor affecting its binding to other TSPANs.

The number and approximate molecular weight of the *E. histolytica* TEM protein complexes seem to vary based on our BN-PAGE analysis of organellar membrane fractions from TSPAN4-HA, TSPAN12-HA, TSPAN13-HA, and TBP55-HA expressing strains (Fig 4). Multiple TEMs were identified in TSPAN12-HA and TBP55-HA expressing stains by anti-HA antibody. The different sizes may be due to different compositions and stoichiometric ratios between TSPANs and other interacting partners. Meanwhile, the absence of a complex in TSPAN13-HA from our BN-PAGE analysis is consistent with the result of MS analysis that TSPAN13-HA was unable to reversely pull-down TSPAN4 and TSPAN12 (S3 Table). This result may indicate that overexpression of TSPAN13, or the epitope tag positioned at the carboxyl terminus, may have a suppressive effect on its complex formation with TSPAN4, TSPAN12 and TBP55, while its binding with *Eh*interaptin seemed unaffected.

In humans, TSPANs play crucial biological roles via their interactions with different single transmembrane-domain binding partners such as integrin. Integrins are transmembrane receptors involved in cell adhesion and communication between the cell and its external environment. They are heterodimeric proteins, composed of two subunits (α and β), which work together to bind to extracellular matrix (ECM) proteins or cell surface molecules [30]. The TSPAN CD63 interacts with various integrins including β1 and β3 integrin in primary endothelial cells [34,35]. CD63 links vascular endothelial growth factor receptor 2 (VEGFR2) to β1 integrins in endothelial cells, thus allowing endothelial cells to respond to extracellular matrix-immobilized angiogenic VEGF [36]. Similarly, CD9 interacts with various integrins including β1 and β3 [12]. Specifically, CD9 was found to link JAM-A, a cell adhesion receptor, to αvβ3 integrin to form a ternary JAM-A–CD9–αvβ3 integrin complex. This complex formation mediates basic fibroblast growth factor-specific angiogenic signaling [37]. Here, we found that *E. histolytica* TSPANs also form diverse complexes and bind with various partners. TBP55 (EHI_001100) was detected in TSPAN4-HA, TSPAN12-HA, and TSPAN13-HA co-IP proteomes in a comparatively high QV, which suggests that TBP55 is a very important binding partner of TSPAN4, TSPAN12, and TSPAN13 (Table 3). TBP55 possesses a signal peptide at the amino terminus and a transmembrane domain at the carboxyl terminus (S5 Fig) and was previously considered as a mitosomal membrane protein [38]. In the Uniprot database, it is annotated as a von Willebrand factor type A (VWFA)-domain containing protein. VWFA proteins share a common feature of forming multiprotein complexes [39], which is also exhibited by TBP55. Interestingly, TBP55 was also detected to be expressed significantly higher in non-virulent strain than virulent strain of *E. histolytica* [40], suggesting that it may also play a role in the suppression of the parasite’s virulence. The identification of short chain dehydrogenase, steroid 5-alpha reductase, and β-keto acyl reductase implies that TBP55 might play a role in the regulation of fatty acid elongation. Furthermore, the presence of vesicular membrane-associated proteins such as coatomer subunit gamma (EHI_040700) and coated vesicle membrane protein (EHI_096210) suggests that TBP55 could also be involved in vesicular transport between the endoplasmic reticulum (ER) and the cis-Golgi network.

**Table 3.** Summarization of the interactions between major binding partners of TSPAN4 based on the MS analysis results of HA-tagged TBP55 co-immunoprecipitation.

| Bait proteins | Trial | QV (TSPAN4 vs Mock-HA) |  | QV (TSPAN12 vs Mock-HA) |  | QV (TSPAN13 vs Mock-HA) |  | QV (TBP55 vs Mock-HA) |  | QV (*Eh*interaptin vs Mock-HA) |  |
| --- | --- | --- | --- | --- | --- | --- | --- | --- | --- | --- | --- |
| TSPAN4<br>(EHI_075690) | 1 | 8.7 | 0 | 13.1 | 0 | 24.0 | 0 | 50.1 | 0 | 10.9 | 0 |
|  | 2 | 6.1 | 0 | 13.8 | 0 | 4.6 | 0 | 55.3 | 1.1 | 1.5 | 0 |
|  | 3 | 28.5 | 0 | 13.8 | 0 | 28.5 | 0 | 210.7 | 0 | 7.9 | 0 |
| TSPAN12<br>(EHI_091490) | 1 | 3.1 | 0 | 4.9 | 0 | 2.5 | 0 | 8.0 | 0.4 | 4.3 | 1.1 |
|  | 2 | 9.8 | 0 | 28.2 | 0 | 17.4 | 0 | 115.0 | 0 | 21.7 | 0 |
|  | 3 | 3.1 | 0 | 14.1 | 0 | 14.1 | 0 | 73.6 | 0 | 0 | 0 |
| TSPAN13<br>(EHI_107790) | 1 | 0 | 0 | 0.4 | 0 | 17.1 | 0 | 0.4 | 0 | 22.7 | 0 |
|  | 2 | 0 | 0 | 0 | 0 | 18.7 | 0 | 1.8 | 0 | 27.0 | 0 |
|  | 3 | 0 | 0 | 0 | 0 | 9.2 | 0 | 4.9 | 0 | 43.6 | 9.5 |
| TBP55<br>(EHI_001100) | 1 | 0 | 0 | 21.7 | 0 | 96.3 | 0 | 85.1 | 0 | 0 | 0 |
|  | 2 | 16.2 | 0 | 11.3 | 0 | 67.5 | 0 | 529.1 | 2.6 | 0 | 0 |
|  | 3 | 8.9 | 0 | 20.0 | 0 | 24.5 | 0 | 189.2 | 0 | 0 | 0 |
| *Eh*interaptin<br>(EHI_148910) | 1 | 0 | 0 | 0 | 0 | 0 | 0 | 0 | 0 | 126.0 | 1.9 |
|  | 2 | 0 | 0 | 0 | 0 | 0 | 0 | 0 | 0 | 44.9 | 1.9 |

The detection of numerous nuclear proteins in our various co-IP experiments using different TSPANs as baits suggests that amebic TSPANs also play a role in nuclear processes. In humans, TSPANs have been implicated in nuclear functions with an example of Tspan8, which was found to carry cholesterol entering the nucleus [41]. Our co-IP MS analysis showed that *Eh*interaptin was consistently identified from transformants expressing TSPAN4-HA, TSPAN12-HA, TSPAN13-HA (S1-S3 Tables). On the other hand, immunoprecipitation of HA-tagged *Eh*interaptin from the corresponding overexpressing transformant failed to pull down TSPAN4, TSPAN12, and TSPAN13, which may indicate a transient nature of their binding (S4 Table). Interaptin was investigated in a previous research in the free-living amoebae *Dictyostelium discoideum* [42]. *Dd*Interaptin (accession number:AF057019) was shown to localized in vesicles of the ER. Both *Dd*interaptin and *Eh*interaptin contain multiple coiled-coil structures based on MARCOIL toolkit prediction (S7A Fig). While canonical *Dd*interaptin possesses an actin-binding domain (ABD) near the amino terminus and a transmembrane helix near the carboxyl terminus, *Eh*interaptin possess neither. Internal repeats in the coiled-coil rod domain are not found in *Eh*interaptin, but when aligned, the putative tyrosine phosphorylation site ‘KKVIDERY’ of *Dd*interaptin has a similar ‘KKRMEELY’ motif in *E. histolytica* (S7B Fig). Besides the annotation as interaptin, this protein was predicted to contain a structural maintenance of chromosomes (SMC) domain, which contains long coiled-coil structures and utilizes ATP hydrolysis to drive conformational changes in DNA topology. SMC proteins are essential for chromosome condensation, segregation, and DNA repair and play a vital role in ensuring the stability and proper functioning of the genome [43]. The in-silico analysis using NCBI’s conserved domain search (https://www.ncbi.nlm.nih.gov/Structure/cdd/wrpsb.cgi) revealed that *Eh*interaptin contains NLS. Furthermore, IFA of *Eh*interaptin-HA showed obvious nuclear, particularly nuclear periphery localization (Fig 3B, top panel). It was previously indicated that the nucleolus is localized at the nuclear periphery in *E. histolytica* [44]. Consequently, it is plausible that *Eh*interaptin is situated in the nucleolus of *E. histolytica*, given the similarity in the observed signals at the nuclear periphery. In eukaryotes, the SMC complexes are usually built from heterodimers of different SMC subunits that form the basis for cohesin, condensin, and Smc5/6 complexes [45,46]. Since *Eh*interaptin possesses an SMC domain with long coiled-coil structure, it can be one subunit of the SMC complex in *E. histolytica.* Also, there is another SMC domain containing protein EHI_152940, which was pulled down by TSPAN4-HA, TSPAN12-HA, and TSPAN13-HA multiple times (S1-S3 Table). In the two trials of MS analysis of *Eh*interaptin-HA co-IP samples, EHI_152940 was also exclusively pulled down in one out of two trials with a very high QV of 56.2. Above these considerations, the data indicate that *Eh*interaptin and EHI_152940 may be potential candidates of SMC subunits in *E. histolytica.* A previous study of a SMC family protein indicates that the knock-out of the Smchd1 lead to a ∼1.5-fold higher expression of TSPAN 8 (Tspan8) in *Homo sapiens* [47]. This evidence suggests that regulation may exist between TSPANs and SMC-domain containing proteins. The binding between *Eh*interaptin and TSPANs shed light on the transient interactions between SMC complexes and TSPANs in *E. histolytica*.

Gal/GalNAc lectin is a surface protein complex that specifically recognizes and binds to Gal/GalNAc on host and bacteria surface molecules including glycoproteins and glycolipids. This lectin plays a pivotal role in the pathogenicity and virulence of *E. histolytic*a [48]. A family of Hgl (heavy subunit: EHI_042370, EHI_077500, EHI_133900, EHI_012270, EHI_046650), Igl (intermediate subunit: EHI_006980, EHI_065330), and Lgl (light subunit: EHI_049690, EHI_159870, EHI_058330, EHI_148790, EHI_183400, EHI_135690, EHI_027800) were previously identified [49]. In the TSPANs proteomes, specific members of Hgl, Igl, and Lgl isotypes were identified (S1 File). Such interactions of TSPANs with Gal/GalNac lectins imply that TEMs in *E. histolytica* may play a specific role such as adhesion and signal transduction. As the integrin-TSPAN adhesion complex was well established and TSPANs can modulate integrin signaling in humans [30]. The cytoplasmic domain of Hgl apparently contains a β2 integrin motif, which regulates the adherence and virulence of *E. histolytica* [50]. Also, Igl2 (EHI_065330) was identified as a β1 integrin-like fibronectin receptor that assembles a signaling complex similar to those of mammalian cells [51]. In the adhesion assay conducted by utilizing TSPAN overexpressing strains, the adhesion capacity was decreased in these overexpressing strains, compared to mock control (Fig 5C). The possible mechanism can be similar with the TSPAN CD82 down-regulating integrin ɑ6-mediated cell adhesion. When CD82 was overexpressed in Du145 prostate cancer cells, the cell surface expression of integrin α6 decreased and the cellular morphogenesis process on matrigel was abolished [52].

Cysteine proteases have been recognized as key contributors to the pathogenicity of numerous infectious organisms [53]. In recent times, several molecules crucial to the pathogenicity of *E. histolytica* have been identified and molecularly characterized [54]. All these molecules, including cysteine proteases, were discovered in both *E. histolytica* and *E. dispar*, casting doubt on their role in *Entamoeba* pathogenicity. One previous research of two major cysteine proteases uniquely present in *E. histolytica* implies that these enzymes are indeed instrumental in the amoeba’s pathogenicity [55]. By CP secretion assay, we showed that trophozoites of *tspan12*gs and *tspan13*gs secrete less CPs into the extracellular environment (Fig 5D). In a previous research, cysteine protease binding protein 1 (CPBF1, EHI_164800) oversaw the maturation and trafficking of *Eh*CP-A5 (EHI_168240) which is one of the major cysteine proteases in *E. histolytica* and responsible for its pathogenicity [56]. The gene silencing of *CPBF1* led to the absence of intracellular mature CP-A5 and enhanced CP-A5 mis-secretion extracellularly [57]. This outcome implies the possibility of an alternative route, aside from lysosome exocytosis, for the transportation of mature CP5 out of the cell, which appears to be compensatorily enhanced when CPBF1 was silenced. The decrease in CP extracellular secretion by *tspan12gs* and *tspan13*gs suggests that these TSPANs are involved in the regulation of *Eh*CPs secretion, plausibly instrumental to the alternative pathway (Fig 6). Additionally, various *Eh*CPBFs and *Eh*CPs were detected in TSPANs’ proteome especially CPBF1 (S5 Table). TSPANs were featured in exosome cargo selection in other organisms[10,58,59]. Thus, it is plausible that TSPAN12 and TSPAN13 are engaged in cargo selection for *Eh*CPs in the ER or endosomes, packaging them into exosomes, transported via multivesicular bodies, and released extracellularly (Fig 6A). Although the detailed mechanism of the maturation and traffic of CP-A5 when CPBF1 was silenced remained unknown, the evidence strongly suggests that the alternative pathway plays an important role in transporting mature CP-A5 into the extracellular environment. Overall, our hypothesis suggests there are at least two pathways for secretion of CPs in *E. histolytica* (Fig 6B and 6C). One of the pathways is through exosome release, where the CP receptor deliver *Eh*CPs into multi-vesicular bodies (MVBs). These CPs are bound and sorted by TSPANs as exosomal cargo followed by their eventual release in these membrane-bound extracellular vesicles. The other pathway involves the default secretory pathway by transport of *Eh*CPs via endosomes and lysosomes. In contrast to the MVB/exosome pathway, *Eh*CPs are directly excytosed to the extracellular milieu by lysosomal fusion to the plasma membrane (Fig 6A).

**Fig 6.**
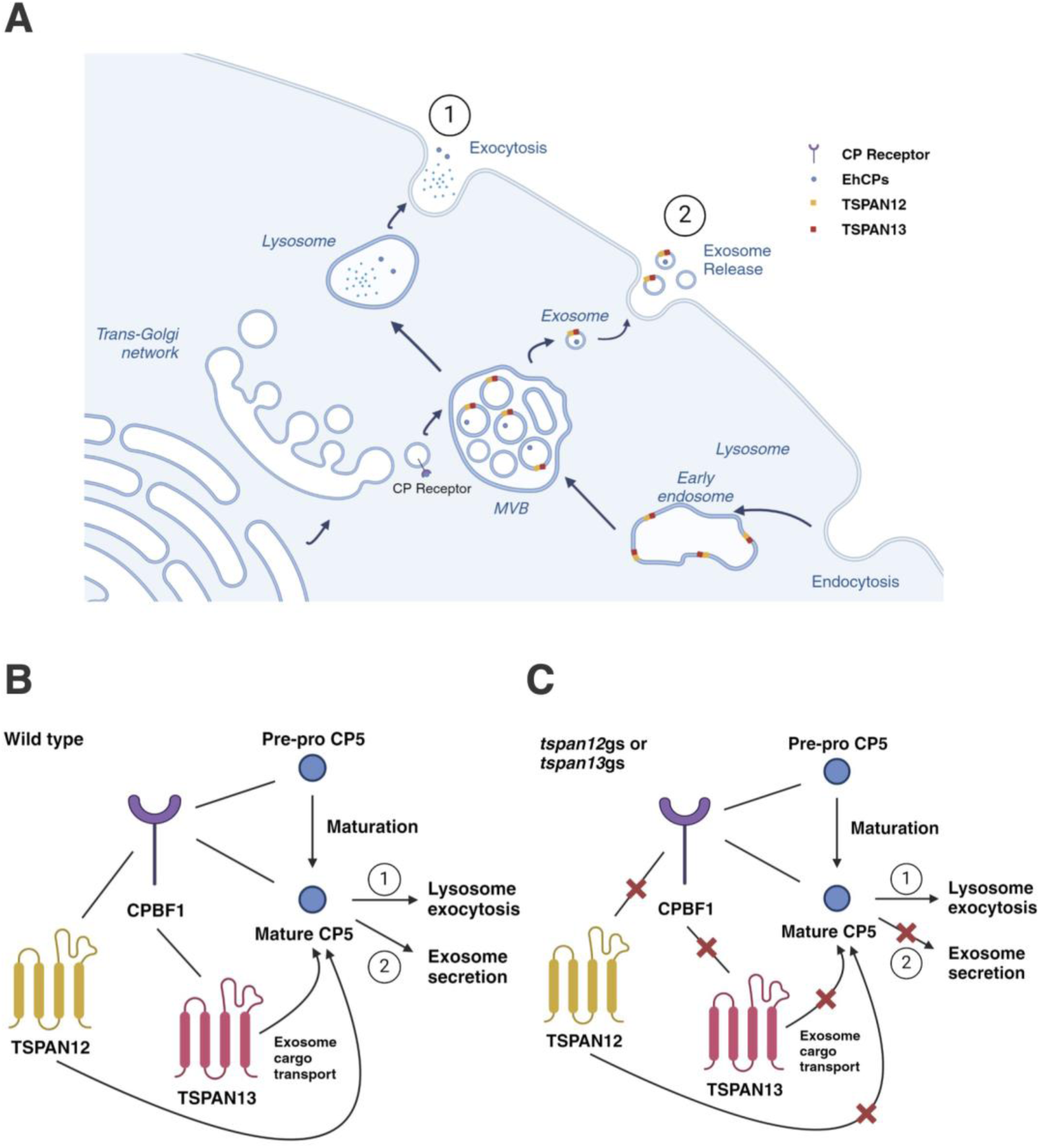
Possible models of regulation of *Eh*CP secretion by *Eh*TSPANs. TSPAN12 and TSPAN13 together with CPBF1 form two pathways of secretion of *Eh*CP-A5. (A) A diagram of the vesicular pathway of global CP secretion in *E. histolytica.* In wild type cells (B), *Eh*CP-A5 is released to the extracellular milieu by both exocytosis and exosome release. When either *tspan12* or *tspan13* is silenced (C), *Eh*CP-A5 is mostly secreted by exocytosis. Created with BioRender.com.

## Materials and methods

### Organisms, cultivation, and reagents

Trophozoites of *E. histolytica* clonal strains HM-1: IMSS cl6 and G3 (Diamond et al., 1972) were cultured axenically in 6 ml screw-capped Pyrex glass tubes in Diamond’s BI-S-33 (BIS) medium (Diamond et al., 1978) and at 35.5°C. The anti-HA 16B12 mouse monoclonal antibody was purchased from Biolegend (San Diego, USA). Lipofectamine, PLUS reagent, and geneticin (G418) were purchased from Invitrogen. CellTracker Green, Orange, and Blue were purchased from Thermo Fisher Scientific (Massachusetts, USA). Restriction enzymes and DNA modifying enzymes were purchased from New England Biolabs (Massachusetts, USA) unless otherwise mentioned. Luria-Bertani (LB) medium was purchased from BD Difco (New Jersey, USA). Other common reagents were purchased from Wako Pure Chemical (Tokyo, Japan), unless otherwise stated.

### Establishment of *E. histolytica* transformants

To construct respective plasmids to express TSPAN4 (EHI_075690), TSPAN12 (EHI_091490), TSPAN13 (EHI_107790), TBP55 (EHI_001100) and *Eh*interaptin (EHI_148910) fused with HA tag at the carboxy terminus, DNA fragments corresponding to coding regions were amplified by polymerase chain reaction (PCR) from *E. histolytica* cDNA using certain designed primers (See Supplementary file 2). The PCR-amplified fragments, and vector plasmids were digested with either *Xho*I and *Xma*I (for N-terminal tagging) or *Bgl*II (for C-terminal tagging), respectively. Amplified fragments were ligated to pEhExHA vector, respectively [60,61]. Ligation between PCR-amplified fragments and vectors were conducted to produce pEhExTSPAN4-HA, pEhExTSPAN12-HA, pEhExTSPAN13-HA, pEhExTBP55-HA, and pEhEx*Eh*interaptin-HA. For G3 strain based antisense small RNA-mediated transcriptional silencing of *tspan4*, *tspan12*, *tspan13*, *tbp55* and psAP2-Gunma vector genes, a fragment measuring 420 base pairs from the gene’s protein-coding section, specifically from the protein’s amino-terminal end, was amplified using PCR from cDNA, this process utilized sense and antisense oligonucleotides that included *Stu*I and *SacI* restriction sites [31,62–64]. Plasmids generated from pEhExHA vector were introduced into the trophozoites of *E. histolytica* HM-1: IMSS cl6 strain, whereas plasmids constructed from the psAP2-Gunma vector was introduced into G3 strain by lipofection as described previously [65]. Transformants were initially selected in the presence of 1 μg/ml G418 until the drug concentration was gradually increased to 50 μg/ml for pEhExTSPAN12-HA, pEhExTSPAN13-HA, and 10 μg/ml for remaining strains. Reverse transcriptase PCR was performed to check mRNA levels of *tspan4*, *tspan12*, *tspan13*, *tbp55* gene silenced strains, total RNA was extracted from trophozoites of gene silenced and control strains that were cultivated in the logarithmic phase using TRIZOL reagent (Life Technologies, California, USA). Approximately 1 µg of DNase treated total RNA was used for cDNA synthesis using Superscript III First-Strand Synthesis System (Thermo Fisher Scientific, Massachusetts, USA) with reverse transcriptase and oligo (dT) primer according to the manufacturer’s protocol. ExTaq PCR system was used to amplify DNA from the cDNA template using the primer set written in S2 File.

### Indirect immunofluorescence assay (IFA)

Approximately 5×10^3^ trophozoites in 50 µl BIS were transferred to a 8 mm round well on a slide glass (Matsunami Glass Ind, Osaka, Japan). After 10 minutes incubation in an anaerobic chamber at 35.5 °C, removing the medium, cells were fixed with PBS containing 3.7% paraformaldehyde at room temperature for 10 minutes, after fixation, wash the cells with 1x PBS for three times, and subsequently permeabilized with PBS containing 0.2% saponin and 1% bovine serum albumin (BSA) for 10 min each at room temperature. The cells were then reacted with anti-HA mouse monoclonal antibody (1:500), anti-HA rabbit monoclonal antibody (1:200), Hgl 3F4 mouse monoclonal antibody (1: 300) [66], anti-Vps26 rabbit polyclonal antiserum (1:500) [29], or anti-BiP polyclonal antiserum (1:800) [67]. for 1 hour at room temperature. Then the sample was reacted with Alexa Fluor-488 conjugated anti-mouse IgG (1:1000) antibody, Alexa Fluor-568 conjugated anti-rabbit IgG (1:1000) antibody (ThermoFisher, Massachusetts, USA) or Hoechst 33342 (diluted into 1 µg/ml) (ThermoFisher, Massachusetts, USA). The images were then captured using LSM 780 confocal microscope and analyzed by ZEN software (Carl-Zeiss, Oberkochen, Germany).

### Immunoblot analysis

*Entamoeba histolytica* trophozoites expressing TSPAN4-HA grown in the exponential phase were harvested and washed three times with phosphate buffer saline (PBS). After resuspension in lysis buffer (150 mM NaCl, 50 mM Tris-HCl, pH 7.5, 0.1% Triton-X 100, 0.5 mg/ml E-64, and 1x cOmplete Mini protease inhibitor cocktail (Roche, Mannheim, Germany)), the trophozoites were incubated on ice for 30 min, followed by centrifugation at 500 x g for 5 min. Around 30 µg of whole cell lysate were separated on 5-20% SDS-PAGE and followed by electro transferring onto nitrocellulose membranes. The membranes were incubated in 5% dried skim milk in Tris-Buffered Saline and Tween-20 (TBST; 50 mM Tris-HCl, pH 8.0, 150 mM NaCl, and 0.05% Tween-20) for 30 min at room temperature to block the non-specific binding sites on the membrane. The blots underwent reaction with one of the following primary antibodies, appropriately diluted in TBST as follows: the anti-HA 16B12 monoclonal mouse antibody at a 1:1,000 dilution, and the anti-CS1 rabbit polyclonal antiserum [68] at a dilution of 1:1,000, all at a temperature of 4°C overnight. Following this, the membranes were washed thrice with TBST and subsequently subjected to further reaction with horseradish peroxidase (HRP)-conjugated anti-mouse or anti-rabbit IgG antiserum, both diluted at a ratio of 1:10,000 and carried out at room temperature for 1 hour. After washing with TBST thrice, the specific proteins were visualized using a chemiluminescence HRP Substrate system (Millipore, Massachusetts, USA) and LAS 4000 (Fujifilm Life Science, Cambridge, USA) or ChemiDoc Imaging System (Bio-Rad, USA) as per the manufacturer’s protocol.

### Co-immunoprecipitation (Co-IP)

Approximately 1×10^6^ trophozoites of the amoeba transformants were cultured in a 10-cm dish using BIS medium under a low oxygen environment provided by Anaerocult (Merck) and incubated at 35.5 °C for 48 hours. Post medium removal, the amoebae were gently dislodged using cold PBS and chilled on ice for 10 minutes. Following three PBS washes, trophozoites were fixed with DSP solution (Thermo Fisher, Massachusetts, USA). The cells, after a couple of PBS rinses, were lysed in 800 µl of lysis buffer. Post debris clearance by centrifugation at 16,000 *g* at 4 °C for 5 minutes, the supernatants were combined with Protein G Sepharose beads (GE Healthcare, Illinois, USA) and incubated at 4 °C for an hour. The supernatants, after a brief centrifugation, were transferred to new microtubes containing anti-HA agarose bead-conjugated monoclonal antibodies (Sigma Aldrich) and incubated with gentle mixing at 4°C for 3.5 hours, followed by centrifugation to remove non-binding proteins. The beads underwent three lysis buffer washes and were then incubated in lysis buffer with HA peptide at a 0.2 mg/ml concentration overnight at 4°C to elute the bound proteins.

### Mass spectrometry

Protein sequencing by mass spectrometry was performed by the Mass Spectrometry and Protemics Core, Johns Hopkins University, Maryland, USA. Each lyophilized protein sample was resuspended in 40µl of 20mM triethylammonium bicarbonate (TEAB), pH 8.0, and reduced with 5µl of 50mM dithiothreitol at 60°C for an hour. After cooling, samples were alkylated with 5µl of 100mM chloroacetamide for 15 minutes in darkness. They were then diluted with 400µl of 9M urea and processed using a 30 kD MWCO spin filter pre-washed with distilled water. Post-washing with urea and TEAB, each sample was treated with 400 ng of enzyme in 300 µl of TEAB and digested overnight at 37°C. The following day, digested peptides were collected post-centrifugation, the filters were rinsed, and the flow-through was combined with the peptides. Peptides were acidified, desalted using an Oasis HLB microelution plate, dried in a speedvac, and reconstituted in 2% acetonitrile with 0.1% formic acid for analysis by a Q-Exactive Plus Hybrid-Quadrupole-Orbitrap mass spectrometer.

### Organellar membrane fractionation

Subcellular fractionation was performed as previously described [69,70], with a few modifications. Six 10-cm dishes of confluent trophozoite cultures in 15 ml BIS medium, for certain transformants, were prepared. After two days of incubation in an anaerobic chamber at 35.5 °C, the medium was removed, and 6 ml of cold PBS were added to the dishes. Cells were harvested into 50 ml centrifuge tubes and were centrifuged at 800 *g* for 3 minutes. Pellets were washed 3 times with 1 ml PBS. The wet pellet was weighed and added with the homogenization buffer (SM buffer; 0.25 M sucrose, 10 mM MOPS-KOH, pH 7.2, 0.5 mg/ml E-64, and 1x cOmplete Mini protease inhibitor cocktail) and then homogenized using a glass homogenizer. The cells were homogenized until approximately 20% of the population of trophozoites remained intact. After centrifugation at 5000 *g* for 10 minutes, the supernatant was collected and subjected to fractionation by ultracentrifugation at 100,000 *g* for 60 minutes. The supernatant was collected as the cytosolic fraction. The remaining pellet was resuspended with PBS and utilized in second ultracentrifugation at 100,000 *g* for 60 minutes at 4 °C. After centrifugation, the supernatant (cytosolic fraction) was collected, and the pellet (organellar fraction) was resuspended with 100 µl PBS. The mild detergent digitonin was added to samples for solubilization on ice for 30 minutes. The samples were centrifuged at 16,000 *g* for 30 minutes, and then the supernatant was collected as it contained solubilized organellar proteins.

### Blue Native-PAGE

Blue Native-PAGE (BN-Page) was conducted following the protocol from NativePAGE™ Novex® Bis-Tris Gel System kit (Thermo Fisher Scientific, Massachusetts, USA). Samples separated by membrane fractionation were utilized in BN-Page. The permeabilized samples were added with NativePage buffer (Thermo Fisher Scientific, Massachusetts, USA) and Coomassie G-250 dye. The BN-Page was performed in XCell™ SureLock™ Mini-Cell (Thermo Fisher Scientific, Massachusetts, USA) followed by electro transferring onto nitrocellulose membranes mentioned above.

### Cysteine protease (CP) secretion assay

CP secretion assay was performed following previous paper with a few modification [57]. Approximately 2.5 × 10^4^ amebic transformants were lysed in 50 μl of Transfection medium (Opti-MEM serum reduced medium, pH 6.8, 5.3 mM ascorbic acid, 41.3 mM L-cysteine) by 3 cycles of freezing and thawing, and debris was removed by centrifugation at 14,000 *g* for 5 min. After preincubation of 1 μl of lysate in 74 μl of assay buffer (0.1 M KHPO_4_ pH 6.1, 1 mM EDTA, 2 mM DTT) and 8 μl of supernatant in 68 μl of assay buffer (0.1 M KHPO_4_ pH 6.1, 1 mM EDTA, 2 mM DTT), at room temperature for 10 min, 25 μl of assay buffer containing 2.5 mM benzyloxycarbonyl-L-arginyl-L-arginine-4-methylcoumaryl-7-amide (z-Arg-Arg-MCA, Peptide Institute Inc., Osaka, Japan) was added and fluorescence signal emitted at 505 nm with excitation at 400 nm was recorded for 15 minutes using a SpectraMax Paradigm multimode microplate reader (Molecular Devices, San Jose, CA, USA). z-Arg-Arg-7-amino-4-frifluoromethylcoumarin (AFC) was utilized as a standard. CP activities were expressed in percentage of mock control. The significance of the data was evaluated by unpaired T-test.

### Cell adhesion assay

Transformants were stained with CellTracker Green for 40 minutes at 35.5°C, after washing by PBS, approximately 40,000 cells were incubated to a 96-well glass plate for 40 minutes at 35.5°C. Fluorescence intensity was measured using a SpectraMax Paradigm multimode microplate reader (Molecular Devices, San Jose, CA, USA) at 492 nm/532 nm right after incubation. Fluorescence was re-measured after discarding the medium carefully and one-time washing with warm BI. The percentage of adhesive cells was computed by dividing the final with the initial fluorescence measurements.

## Acknowledgements

We thank all the members of the Nozaki laboratory at the University of Tokyo for the helpful discussion. We thank Seiji Fujimoto and Mihoko Imada for construction of several plasmids and technical support. We also thank William Petri for the anti-Hgl antibody. We also acknowledge Yumiko Saito Nakano and Seiki Kobayashi of the National Institute of Infectious Diseases Japan for providing additional support.

## Supporting information

**S1 Fig. Validation of the co-immunoprecipitation of TSPAN12-HA.** Representative Immunoblot analysis and silver staining of TSPAN12-HA in co-IP experiments. (A) 5 µl of each fraction collected from co-IP experiments were added into SDS-PAGE and immunoblot analysis using anti-HA antiserum. Black arrows indicate TSPAN12-HA expression and red arrows indicate the heavy and light chain of anti-HA antiserum. (B) Silver staining was performed by adding 20 µl of eluted fraction of TSPAN12-HA and HA-mock in the co-IP. The black arrows suggest specific bands in the TSPAN12-HA sample.

**S2 Fig. Validation of the co-immunoprecipitation of TSPAN13-HA.** Representative Immunoblot analysis and silver staining of TSPAN13-HA in co-IP experiments. (A) 5 µl of each fraction collected from co-IP experiments were added into SDS-PAGE and immunoblot analysis using anti-HA antiserum. Black arrows indicate TSPAN13-HA expression and red arrows indicate the heavy and light chain of anti-HA antiserum. (B) Silver staining was performed by adding 30 µl of eluted fraction of TSPAN13-HA and HA-mock in the co-IP. The black arrows suggest specific bands in the TSPAN13-HA sample.

**S3 Fig. Validation of the co-immunoprecipitation of TBP55-HA.** Representative Immunoblot analysis and silver staining of TBP55-HA in co-IP experiments. (A) 5 µl of each fraction collected from co-IP experiments were added into SDS-PAGE and immunoblot analysis using anti-HA antiserum. Black arrows indicate TBP55-HA expression and red arrows indicate the heavy and light chain of anti-HA antiserum. Green arrows indicate truncated TBP55-HA. (B) Silver staining was performed by adding 20 µl of eluted fraction of TBP55-HA and HA-mock in the co-IP. The black arrows suggest specific bands in the TBP55-HA sample.

**S4 Fig. Validation of the co-immunoprecipitation of *Eh*interaptin-HA.** Representative Immunoblot analysis and silver staining of *Eh*interaptin-HA in co-IP experiments. (A) 5 µl of each fraction collected from co-IP experiments were added into SDS-PAGE and immunoblot analysis using anti-HA antiserum. Black arrows indicate *Eh*interaptin-HA expression and red arrows indicate the heavy and light chain of anti-HA antiserum. (B) Silver staining was performed by adding 20 µl of eluted fraction of *Eh*interaptin-HA and HA-mock in the Co-IP. The black arrows suggest specific bands in *Eh*interaptin-HA sample.

**S5 Fig. Domain organization of TSPANs and TBP55 from *Entamoeba histolytica.*** TM (Transmembrane domain), SEL (Small extracellular loop), LEL (Large extracellular loop), SP (Signal peptide), red triangles stand for intracellular domains of tetraspanins.

**S6 Fig. Alignment of TSPANs in *Entamoeba histolytica.*** Multiple amino acids sequence alignment of *Entamoeba histolytica* TSPAN4(EHI_075690), TSPAN12(EHI_091490), TSPAN13(EHI_107790) was established by clustal W algorithm (https://www.genome.jp/tools-bin/clustalw). The highly conserved CCG motifs and the following conserved tryptophan and lysine residues located at large extracellular loops are shown in red font.

**Figure S7. Partial amino acids sequence alignment between Ddinteraptin and Ehinteraptin and Coiled-coil (CC) structure prediction for Ehinteraptin (EHI_148910).** (A) Multiple coiled-coil structures were predicted in Ehinteraptin (EHI_148910) by MARCOIL toolkit (https://toolkit.tuebingen.mpg.de/tools/marcoil) which is a HMM model for coiled-coil domains. (B) Multiple amino acids sequence alignment between Ddinteraptin (AF057019) and Ehinteraptin (EHI_148910) were built by clustal W algorithm. The putative tyrosine phosphorylation site ‘KKVIDERY’ in Ddinteraptin is shown in blue, while the similar motif in Ehinteraptin ‘KKRMEELY’ is shown in red.

**S1 Table. Mass-spectrometry results of HA-tagged TSPAN4 co-immunoprecipitation.** Experimental details referred to Table 1 description but with the bait protein HA-tagged TSPAN4. The list order is sorted by frequency of identification firstly and mean of quantitative value secondly.

**S2 Table. Mass-spectrometry results of HA-tagged TSPAN12 co-immunoprecipitation.** Experimental details referred to Table 1 description but with the bait protein HA-tagged TSPAN12. The list order is sorted by frequency of identification firstly and mean of quantitative value secondly.

**S3 Table. Mass-spectrometry results of HA-tagged TSPAN13 co-immunoprecipitation.** Experimental details referred to Table 1 description but with the bait protein HA-tagged TSPAN13. The list order is sorted by frequency of identification firstly and mean of quantitative value secondly.

**S4 Table. Mass-spectrometry results of HA-tagged *Eh*interaptin co-immunoprecipitation.** Experimental details referred to Table 1 description but with the bait protein HA-tagged *Eh*interaptin. The Co-IP and MS analysis were conducted twice independently. The list order is sorted by frequency of identification firstly and mean of quantitative value secondly.

**S5 Table. *Eh*CPBFs and *Eh*CPs proteins pulled-down by amebic TSPANs and their binding proteins.** *Eh*CPBFs and *Eh*CPs proteins found in TSPANs-HA and TBP55-HA proteome. The upper panel of TSPAN4-HA, TSPAN12-HA, TSPAN13-HA, TBP55-HA indicate the number of independent trials the protein was pulled down by the certain bait protein. The lower panel suggests the mean QV that the protein detected in TSPANs or TBP55, the number inside bracket suggests mean QV that the protein detected in the mock control. ND, no detection.

**S6 Table. Percentage of amino acid identity among TSPAN4, TSPAN12 and TSPAN13.** The alignment was conducted by ClustalW multiple sequence alignment toolkit.

**S1 File. Pull down list for all mass-spectrometry experiments.**

**S2 File. Primer sequences used in experiments.**

